## Supplementary Figure S1 for "ASTROGLIOSIS AND NEUROINFLAMMATION UNDERLIE SCOLIOSIS UPON CILIA DYSFUNCTION"

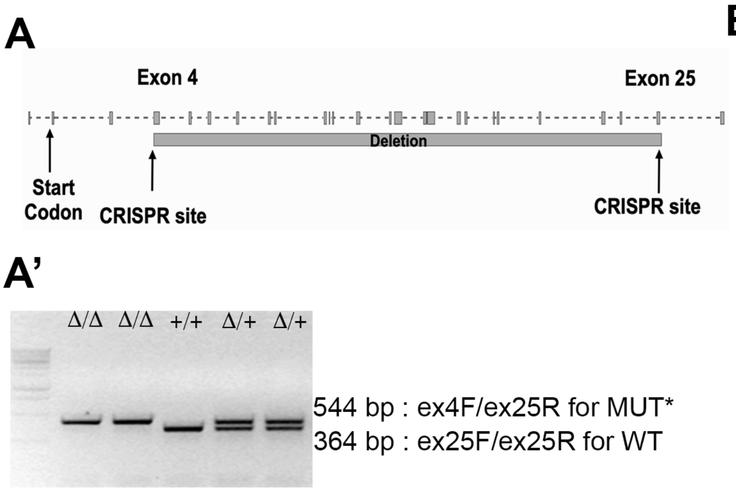


B

| | Heart looping | Kidney cysts | Total nb embryos |
|---|---|---|---|
| wt | 1 | 0 | 900 |
| *rpgrip1l*$^{ex4/ex4}$ | 0 | 0 | 73 |
| *rpgrip1l*$^{\Delta/\Delta}$ | 3 | 0 | 694 |

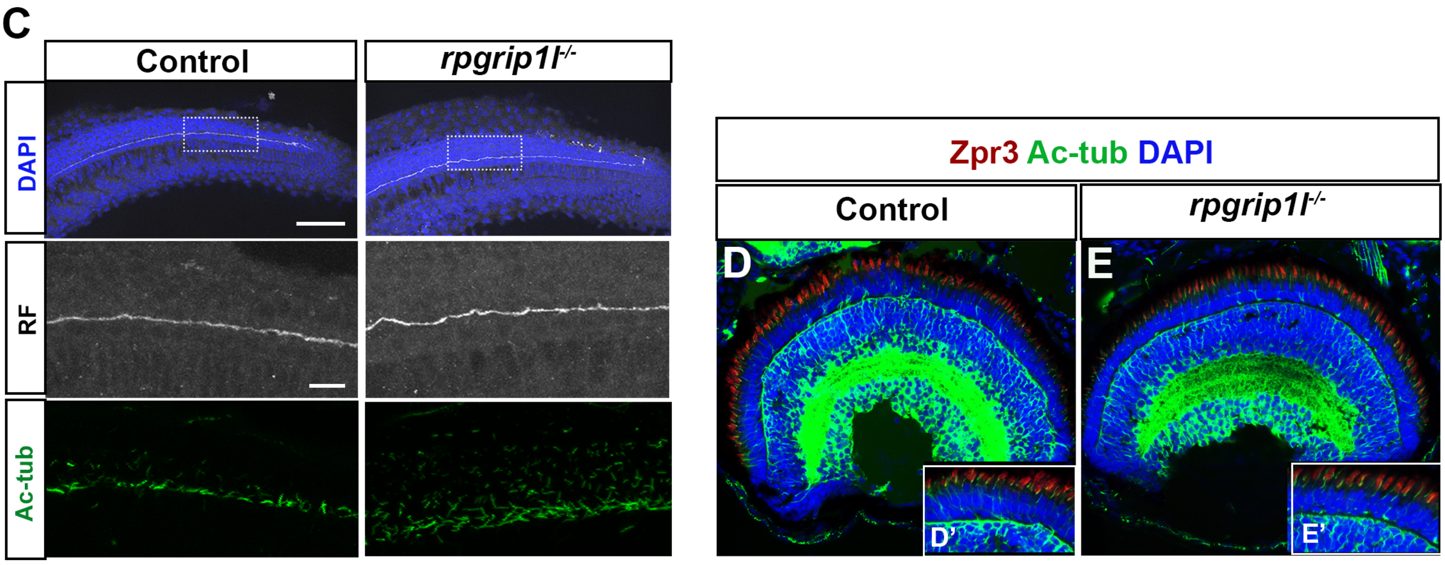


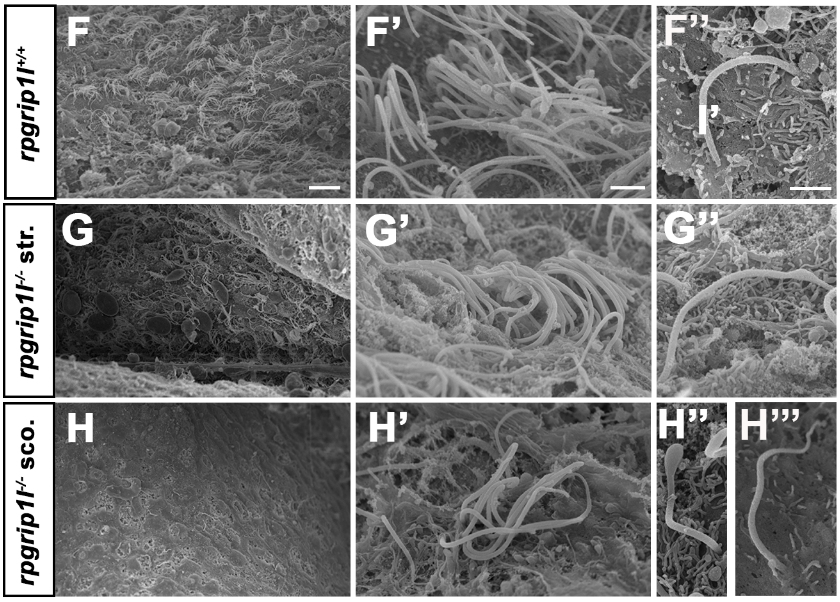


**Supplementary Figure S1**
