## Supplementary figures and images for "ASTROGLIOSIS AND NEUROINFLAMMATION UNDERLIE SCOLIOSIS UPON CILIA DYSFUNCTION"

### Graphical Abstract

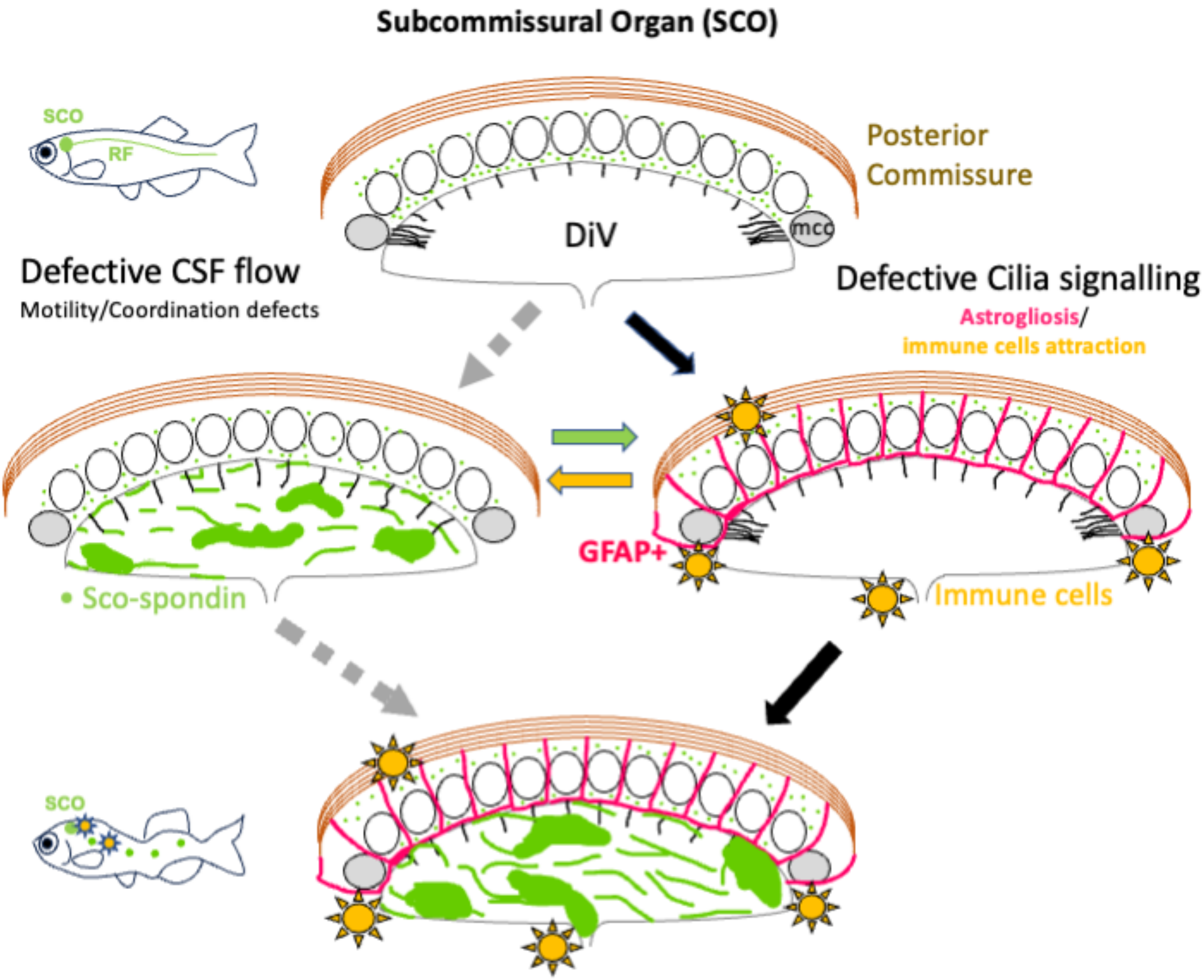

### SSupplementary Figure S3

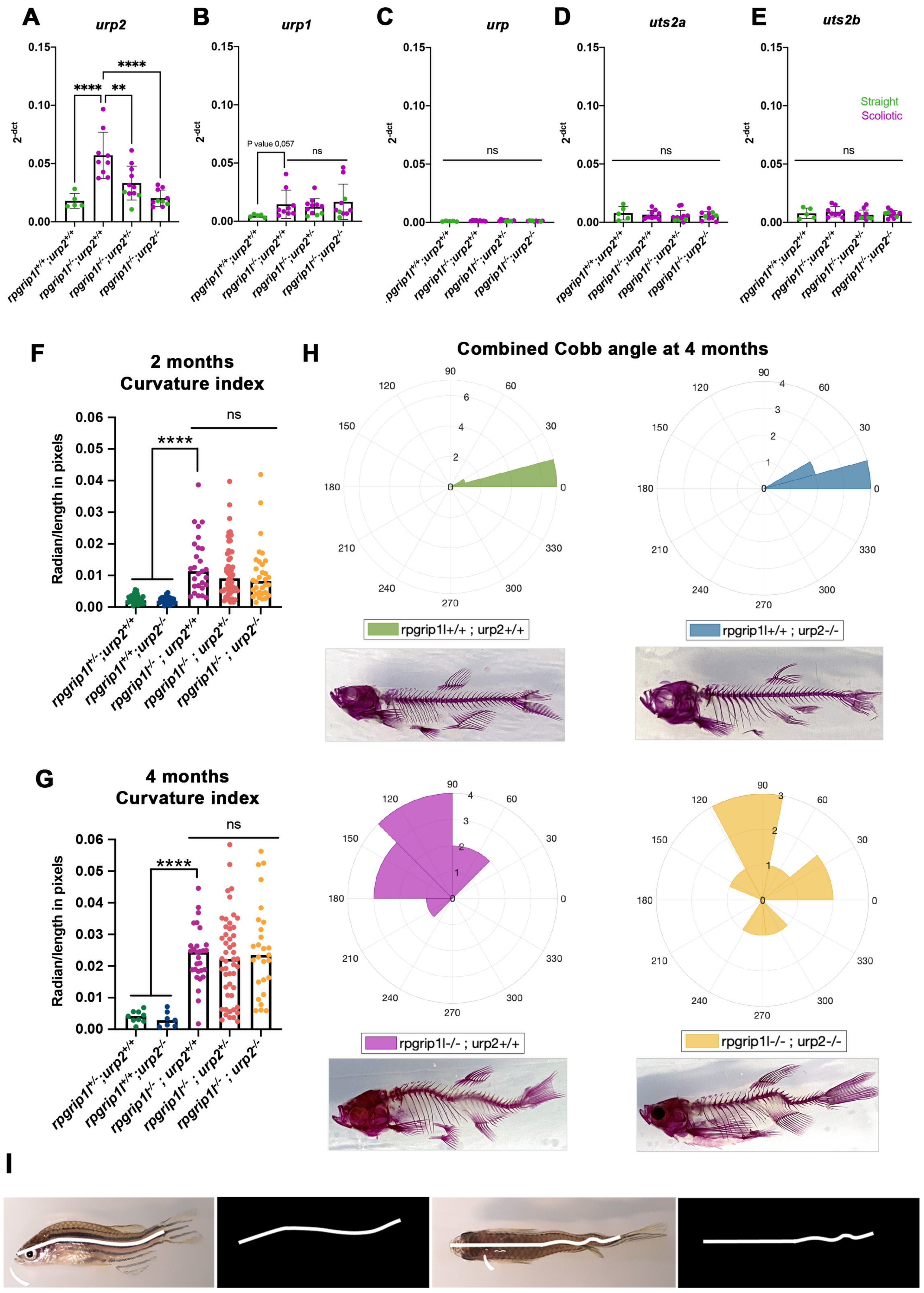


**Supplementary Fig S3**

### Supplementary Figure S2

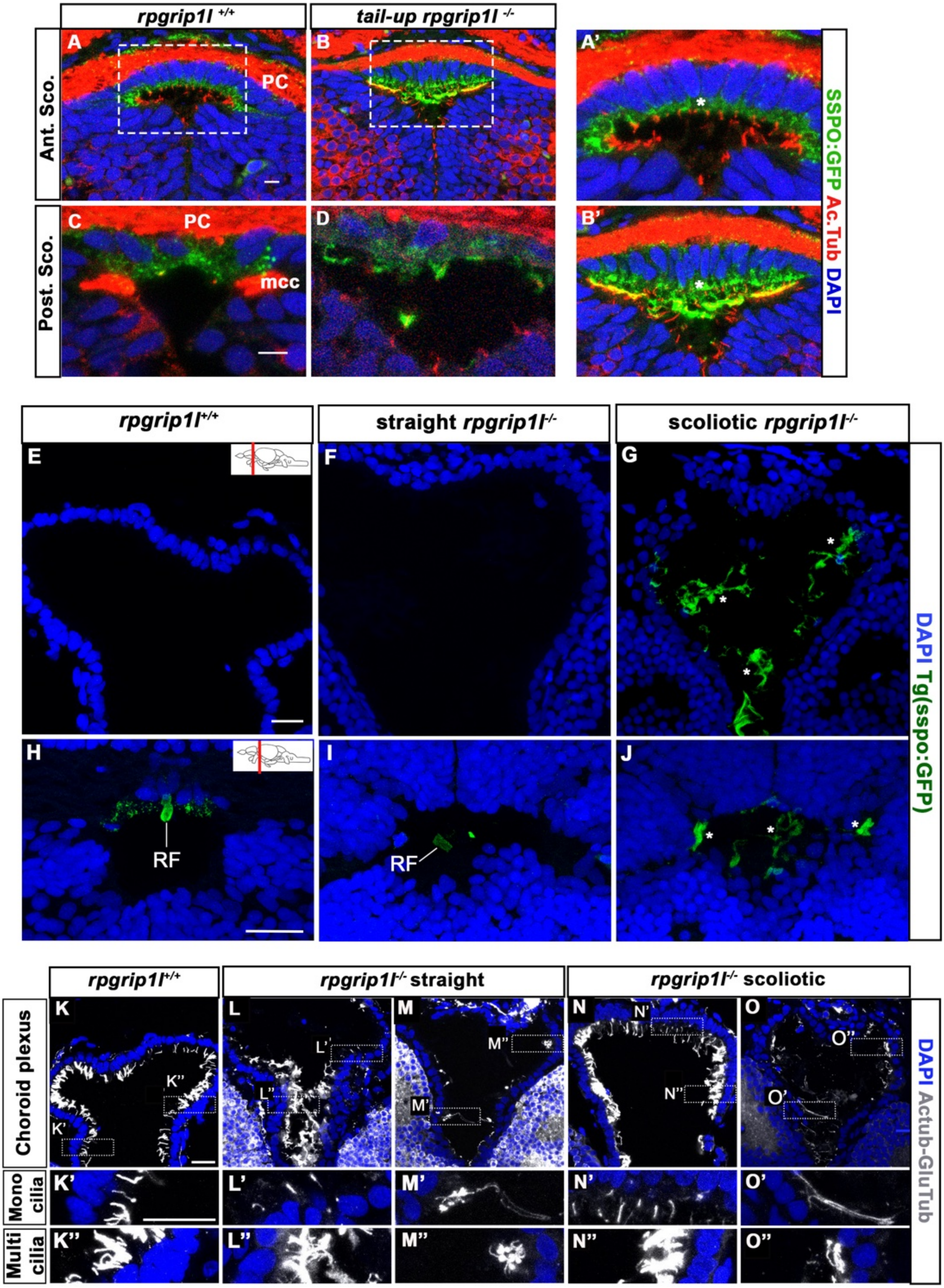


**Supplementary Figure S2**

### Supplementary Figure S5

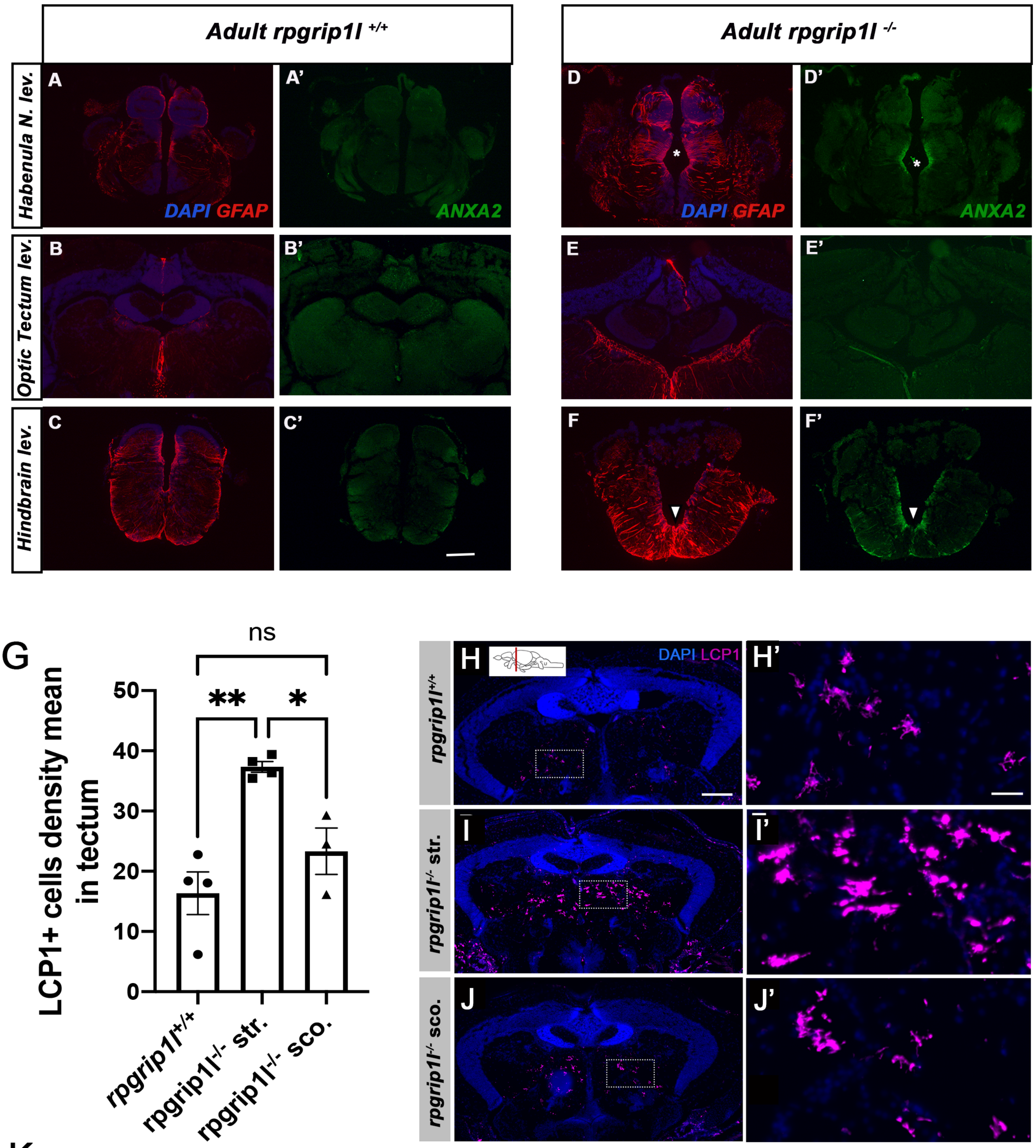


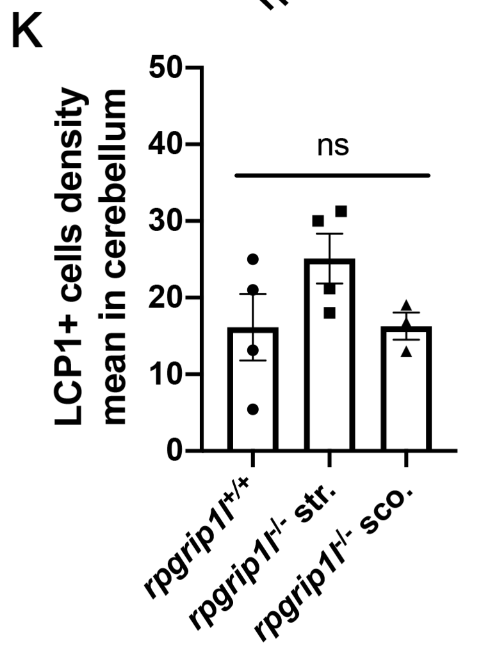


**Supplementary Figure S5**

### Supplementary Figure S6

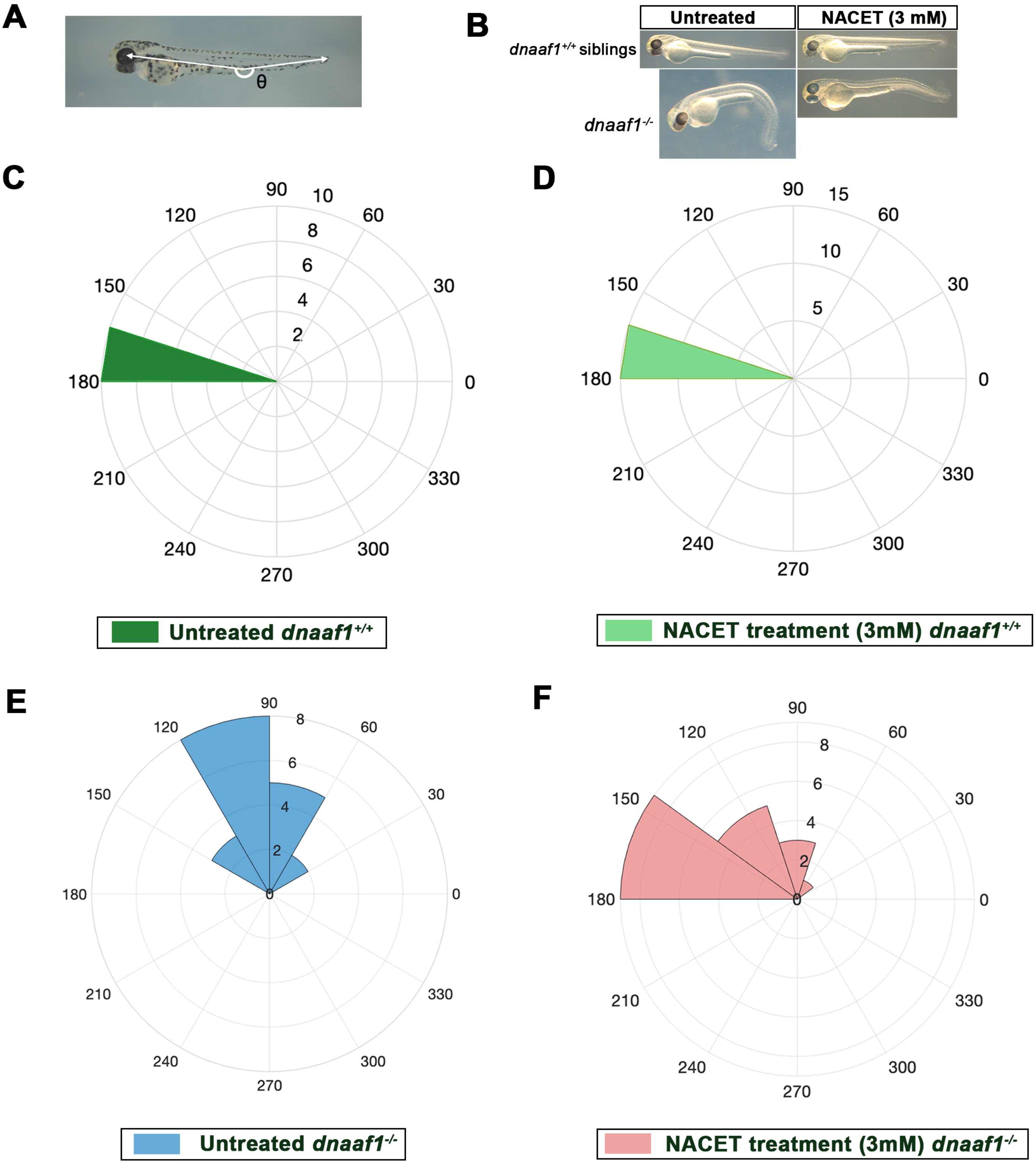


#### Supplementary Figure S6
