## Supplementary Figure S4 for "ASTROGLIOSIS AND NEUROINFLAMMATION UNDERLIE SCOLIOSIS UPON CILIA DYSFUNCTION"

### A: Up-regulated genes in trunk

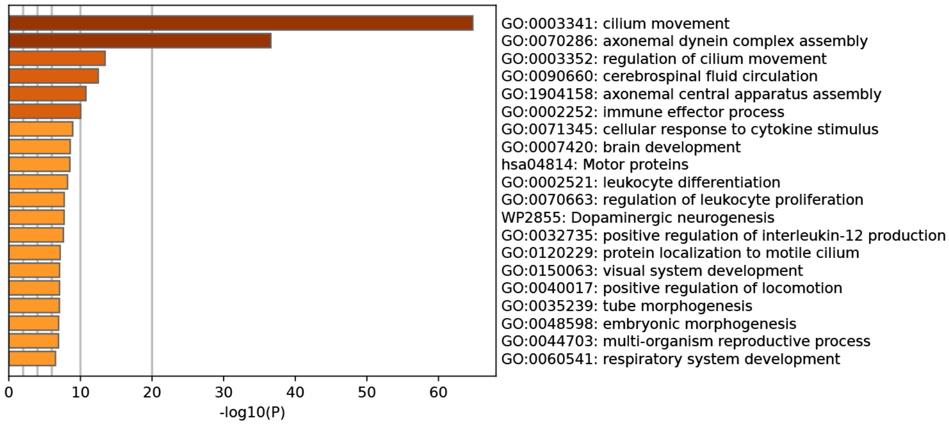


### B: Enriched proteins in adult brain

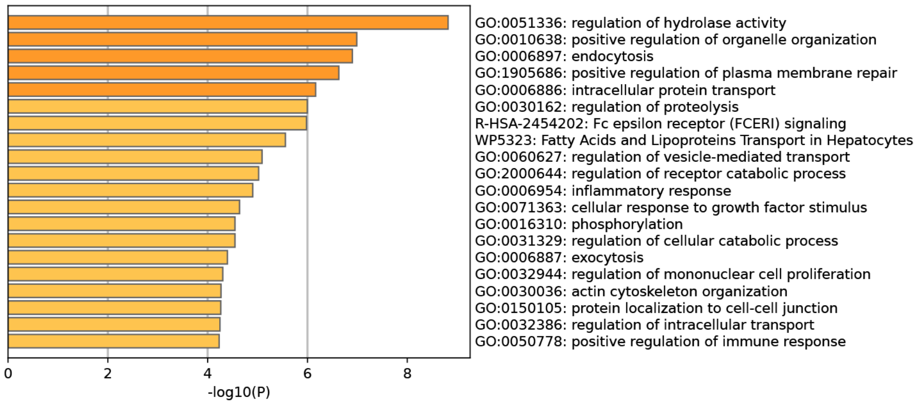


### C: 25 highest P Value

| Protein | FC | P value |
|---|---|---|
| anxa2a | 7,28 | 4,82E-08 |
| echdc2 | 4,97 | 1,99E-06 |
| anxa1a | 6,06 | 9,6856E-06 |
| anxa5b | 2,37 | 3,3303E-05 |
| capsla | 3,72 | 0,00011272 |
| tagln2 | 2,53 | 0,00011315 |
| psph | 1,72 | 0,00011655 |
| c7a | 3,07 | 0,0002431 |
| mapk9 | 2,49 | 0,00029106 |
| vav3b | 2,31 | 0,00036912 |
| serpina7 | 2,46 | 0,0003851 |
| gps1 | 1,92 | 0,00057247 |
| synrg | 2,08 | 0,00066171 |
| c3 like | 3,77 | 0,00068862 |
| tut7 | 2,01 | 0,00102053 |
| gsk3bb | 2,40 | 0,00109597 |
| tmed3 | 3,44 | 0,00148983 |
| mrc1a | 2,10 | 0,00152123 |
| wdr12 | 2,78 | 0,00186884 |
| wdfy2 | 1,97 | 0,00235225 |
| plcd4b | 2,24 | 0,00249521 |
| itchb | 1,81 | 0,00250875 |
| ehd1b | 1,94 | 0,00253596 |
| exoc6 | 1,77 | 0,00298918 |
| stat3 | 5,31 | 0,00319733 |

### D: Activated astrocytes markers

| Hum. Gene | Zf Gene | FC (trunk) | P.adj. Value |
|---|---|---|---|
| **S100B** | s100b | 2,98 | 7,35711E-27 |
| **C4b** | c4b | 5,15 | 1,48224E-21 |
| **GFAP** | gfap | 2,49 | 1,99844E-09 |
| **CTSB** | ctslb | 3,60 | 1,29502E-08 |
| **CAPS** | capsla | 2,67 | 2,31406E-06 |
| **GLAST/SLC1A3** | slc1a3b | 2,68 | 1,46788E-05 |
| **PLCXD3** | plcxd3 | 1,74 | 1,6557E-05 |
| **VIM** | viml | 2,14 | 8,30568E-05 |

**Supplementary Figure S4**
